# Sialylation and fucosylation changes of cytidine monophosphate-Nacetylneuraminic acid hydroxylase (CMAH) and glycoprotein, alpha1,3-galactosyltransferase(GGTA1) knockout pig erythrocyte membranes

**DOI:** 10.1101/2020.08.07.240846

**Authors:** Hak Myong Choe, Zhao-Bo Luo, Mei-Fu Xuan, Biao-Hu Quan, Jin-Dan Kang, Myung Jin Oh, Hyun Joo An, Xi-jun Yin

## Abstract

The recent GGTA1 and CMAH DKO pigs have made it possible to resolve the immune barriers which are duo to xenoantigens on RBC such as αGal and Neu5Gc. Nevertheless, it still requires the detection of glycosylation alternation on the pig RBCs because even the minor changes would be unexpected xenoantigens.

DKO RBC immune reactivity with human serum was assessed by hemagglutination assay. Glycosylation alteration of RBC membranes was characterized by NanoLC-Q-TOF-MS system and lectin blotting assay.

Twelve GGTA1/CMAH DKO piglets were successfully produced. The immunoreactivity with human serum was remarkably reduced in DKO (less than 1:2 dilution), whereas wild type(WT) pigs showed agglutination (the least 1:256 dilution). The MS results showed that DKO increased neutral N-glycans as well as decreased total sialylated N-glycans, especially suggesting significant decrease of di-sialylated N-glycans (P < 0.05). Moreover, lectin blotting assay revealed that DKO pigs reduced the binding signals with AAL, AOL, LCA and SNA and increased the binding signal with MAL.

DKO pigs decreased the expression of total fucosylation and sialylated N-glycans on the erythrocyte membrane. Our findings will support further investigation into DKO pig RBC glycosylation and contribute to uncover the roles of glycan changes for xenotransfusion.

**Summary statement:** To detect glycosylation changes in red blood cells(RBC) of GGTA1/CMAH double knockout(DKO) pigs, comparative analysis of the glycan profiling was done.

## INTRODUCTION

Recent advances in both gene editing engineering and glycobiology have improved largely the research for xenotransfusion. Pig erythrocytes for human xenotransfusion have been attracting more interests of clinical researchers due to having some advantages and sharing many similar characteristics with human erythrocytes(Zhu. 2000; Cooper. 2010; Eckermann. 2004) The pig erythrocytes do not express histocompatibility antigens (i.e., swine leukocyte antigens, SLA), thus reducing immunogenicity and lack of nuclei which would harbor porcine endogenous retroviruses(Oostingh. 2002; Patience. 1998; Blusch. 2002). The inexhaustible pig erythrocytes can replace human erythrocytes for long-term xenotransfusion since easy infection by microorganisms doesn’t make human blood possible to conserve for a long time(Cooper. 2010). The pig is also known as having genetically and structurally similar blood group system with human, and pig erythrocytes with compatible blood type can be used in xenotransfusion to reduce erythrocyte agglutination(Sachs. 1992).

However, pig erythrocytes possess carbohydrate epitopes such as galactose α1,3 galactose (αGal) and N-glycolylneuraminic acid (Neu5Gc), which can cause hyper-acute and acute immune rejection in xenotransfusion(Wang et al. 2014; Jang et al. 2013). The developments of gene editing engineering including zinc finger nucleases (ZFN), transcription activator-like effector nucleases (TALENs) and clustered regularly interspaced short palindromic repeats (CRISPR/Cas9) have made it possible to remove this epitopes(Yang and Wu. 2018).

Some studies reported the investigation into GGTA1 KO and GGTA1/CMAH DKO pig erythrocytes for pig-to-human xenotransfusion with the regard to human antibody-mediated immune reaction of KO pig erythrocytes(Wang et al. 2014; Rouhani et al. 2004). The intestine and pancreas of GGTA1 KO miniature swine revealed that GGTA1 KO pigs increased blood group H type 2 core oligosaccharides and decreased total amount of gangliosides compared to WT(Diswall M et al. 2007). In addition, MS analysis of GGTA1/CMAH DKO pig serum protein showed higher relative amounts of mannosylated, incomplete, and xylosylated N-linked glycans as well as bi- and tri-antennary fucosylated N-linked glycans than human and WT pig serum proteins(Burlak et al. 2013). GGTA1 KO porcine islets reported a slight increase of glycans with α2,3 and α2,6 silaic acid(Shuji et al. 2013). Moreover, GGTA1 KO pig fibroblasts showed significantly higher quantity of fucosylated N-glycans and slightly higher quantity of Neu5Gc than WT pig fibroblasts(Park et al. 2015). However, the previous studies indicated that glycosylation changes had been investigated with pig organs or tissues, but not with erythrocytes.

It is necessary to detect glycosylation changes on glycoproteins and glycolipids in order to identify the roles on the immune rejection because the new glycan pattern can induce and control the immune rejection cascades through the glycan ligand-receptor binding(Brock et al. 2012; Butler et al. 2016; Hryhorowicz et al. 2017). Furthermore, it is known that the glycosylation changes and the immunogenicity of GGTA1 KO or GGTA1/CMAH DKO pigs are different from sample origins since the porcine glycome expression is tissue specific(Jang et al. 2013; Ji et al. 2017; Zhang et al. 2017).

Thus far, comparative analysis of the glycan profiling patterns on organs and tissues from WT and CMAH/GGTA DKO pigs has been widely studied, but little is known about glycosylation changes on DKO pig erythrocyte membrane. Here, we present for the first time experimental data for glycosylation changes on DKO pig RBC membrane through ECC(Extracted Compound Chromatogram)s, CID(collision–induced dissociation) MS/MS and lectin blotting assay, showing the significant decrease of the total fucosylation and sialylated N-glycans.

## RESULTS

### GGTA1 KO fetus production

Porcine ear fibroblasts collected from a donor female mini pig were transfected with the efficient CRISPR/Cas9 plasmids designed to induce the targeted mutation at GGTA1 gene and plasmids encoding enrichment reporters (eGFP and H-2K^k^) by electroporation, and the transfected cells were enriched magnetically. H-2k^k^-positive cells were cultured for two additional days and were used as donor cells in first round SCNT. The GGTA1 mutated embryos generated by the SCNT were transferred to one recipient. Three fetuses were surgically collected on Day 23 of gestation (Figure 1a). The fetuses were analyzed individually with T7E1 assays and sequencing covering the target locus. Mutations at the target locus (GGTA 1 gene) were present in one of the three fetuses. The fetus (F3) carried biallelic mutations and the other two fetuses (F1 and F2) did not show the mutations in GGTA1 gene. The fetus F3 had a 1 bp deletion in both alleles (Figure 1b).

**Figure 1.**
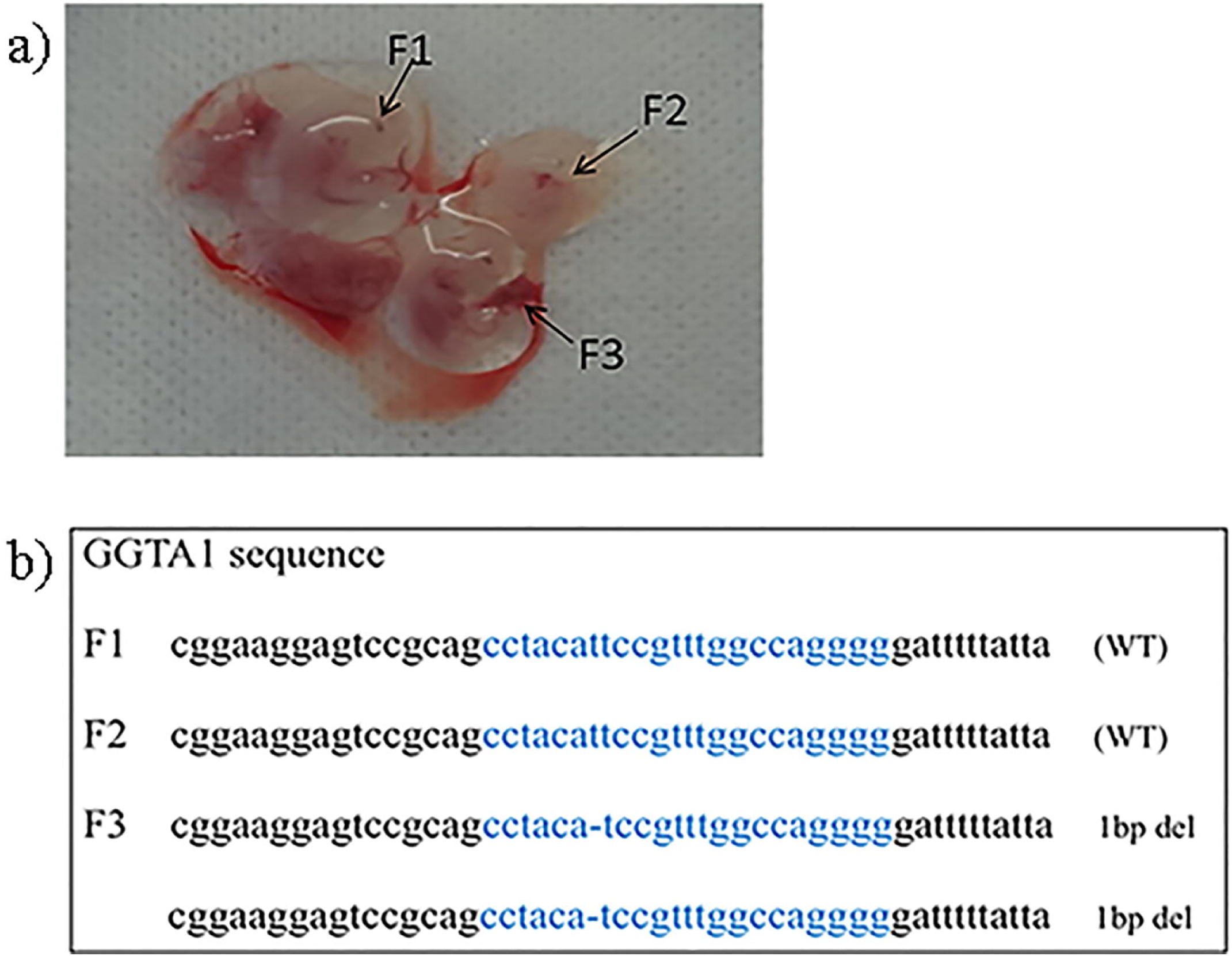
Photographs of the fetuses and GGTA1 sequences after first round SCNT. **(a)** Fetuses (F1,F2,F3) on Day 23 of gestation**. (b)** GGTA1 sequences derived from the harvested fetuses. DNA sequences of the WT and mutant clones. The target sites were highlighted with blue color. The deletion bases were indicated by dashes

### GGTA1/CMAH DKO fetus production

To add CMAH mutation, porcine fetal fibroblasts from the fetus F3 with GGTA1 mutation were collected and transfected with both the CRISPR/Cas9 plasmids and the enrichment reporter plasmids by electroporation. The embryos produced from GGTA1/CMAH mutated donor cells by the second round SCNT were transferred to two recipients and nine fetuses were collected on Day 26 of gestation (Figure 2a). The nine fetuses were analyzed individually with T7E1 assays and sequencing covering the target locus. The homozygous mutations at the target locus (CMAH gene) were detected in fetuses F3-6 and F3-8 of the nine fetuses. Fetus F3-6 had a 8 bp deletion in one allele and a 1 bp insertion in the other allele and fetus F3-8 had a 1 bp insertion in one allele and a 1 bp insertion in the other allele. Fetuses (F3-2, F3-7, F3-9) did not carry mutations and fetuses and (F3-1, F3-3, F3-4) had heterozygous mutations. Fetus F3-5 had homozygous mutatations considered as out of frame (Figure 2b).

**Figure 2.**
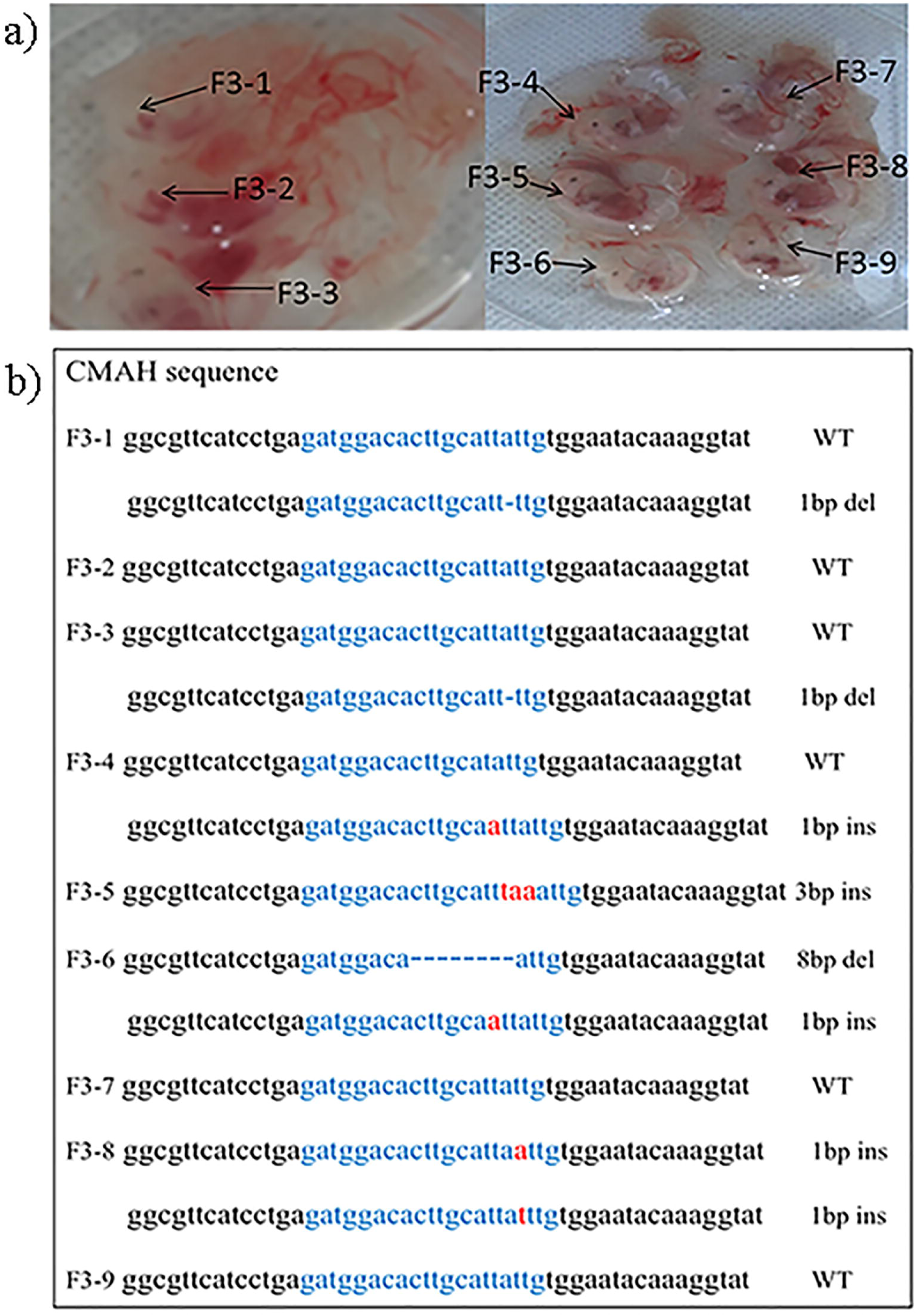
Photographs of the fetuses and CMAH sequences after second round SCNT. **(a)** Fetuses (F3-1,2,3,4,5,6,7,8,9) on Day 26 of gestation. **(b)** CMAH sequences derived from the harvested fetuses. DNA sequences of the WT and mutant clones. The target sites were highlighted with blue color. The deletion bases were indicated by dashes. The inserted bases were indicated by red characters

### GGTA1/CMAH DKO piglet production and phenotype analysis for αGal loss

Fetus F3-6 was selected to produce donor cell lines in this study. GGTA1/CMAH mutated PFFs from the fetus F3-6 were used in third round SCNT and a total of 848 cloned embryos were transferred to four surrogates in estrus (Table I). The twelve piglets were delivered in seven months (Figure 3a). Mutations at the target locus (GGTA1 and CMAH genes) were analyzed and all of them were confirmed with successful mutations in both alleles (Figure 3c). In addition, thirty-three potential off-target sites were investigated, and no off-target events were detected in the live piglets (data not shown).

**TABLE 1.** Results of somatic cell nuclear transfer using double-KO cells

| Recipient | Reconstructed embryos | Pregnancy | Live born |
| --- | --- | --- | --- |
| 1 | 189 | + | 3 |
| 2 | 190 | + | 3 |
| 3 | 263 | + | 2 |
| 4 | 206 | + | 4 |
| Total | 848 |  | 12 |

**Figure 3.**
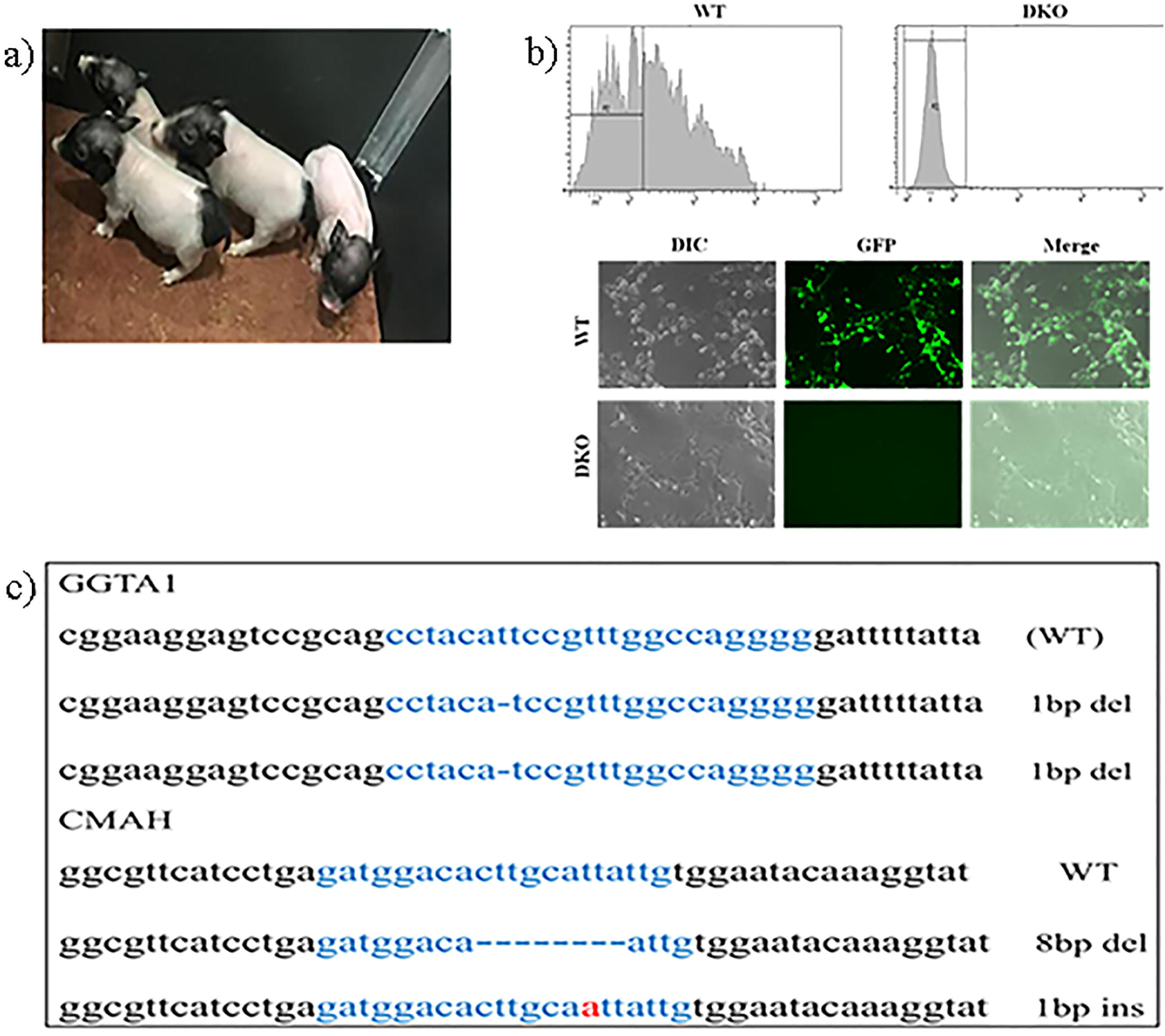
Photographs of viable piglets and GGTA1/CMAH sequences. **(a)** A photograph from the delivered piglets by third round SCNT **(b)** Gal expression on DKO pig skin fibroblasts by FACS analysis and immunofluorescence staining **(c)** GGTA1 and CMAH gene sequences derived from the harvested piglets. DNA sequences of the WT and mutant clones. The target sites were highlighted with blue color. The deletion bases were indicated by dashes. The inserted bases were indicated by red characters

To confirm the phenotype of αGal loss in the produced DKO piglets, the skin fibroblasts were analyzed by FACS analysis and immunofluorescence staining (Figure 3b). In FACS analysis, αGal was absent on DKO pig skin fibroblasts whereas αGal was present on WT. Furthermore, the immunofluorescence staining showed green colored staining in WT, while no staining in DKO.

### Hemagglutination by human serum

Hemagglutination was used to determine total antibody binding reactivity of pooled human directed towards pig erythrocytes. The agglutination image by a 2-fold serial dilution of heat-inactivated human serum was shown (Figure 4). When incubating with human serum (type O), WT erythrocytes agglutinated until 1:256 dilution whereas DKO erythrocytes showed remarkably decreased agglutination with less diluted serum solution than 1:2.

**Figure 4.**
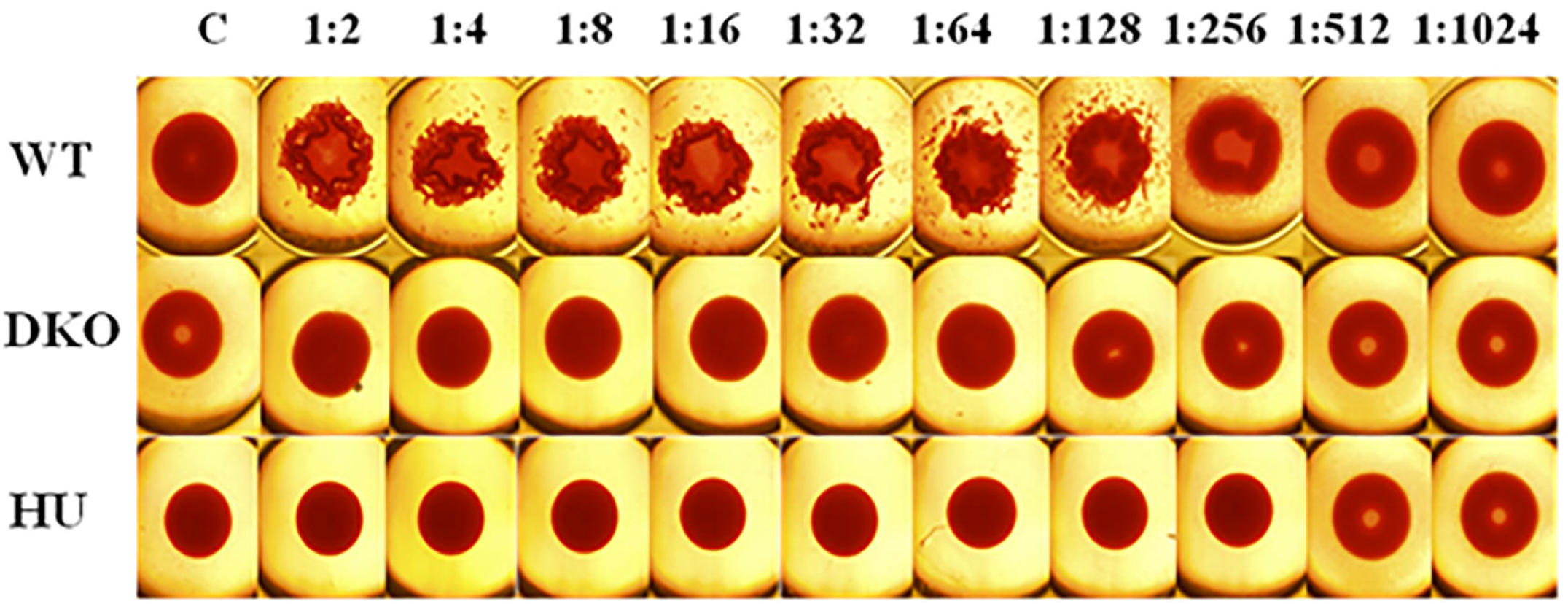
Hemagglutination by human sera. Heat-inactivated human blood type O sera was serially diluted in HBSS and incubated with erythrocytes overnight at room temperature. Untreated erythrocytes were used as negative controls. The 96-well plates were scanned with an epifluorescence microscope. C (negative control); WT (wild type); KO (GGTA1/CMAH double KO); HU (human)

### N-glycan profiling comparison by ECCs and CID MS/MS

In order to explore the sialylated N-glycan profiling, N-glycans released from DKO and WT erythrocyte membranes - were examined by PGC LC/MS of NanoLC-Q-TOF-MS system. The representative extracted compound chromatograms (ECCs) of sialylated N-glycans on WT and DKO RBC membranes were shown in Figure 5a and 5b, respectively. The smaller neutral glycans (high mannose and unsialylated glycans) were eluted first and the larger and more complex glycans were eluted later through PGC column. In particular, sialylated glycans have relatively longer retention in the column than other glycans, so the corresponding RT increases if the degree of sialylation increases. Figure 5a and 5b displayed a direct comparison of individual sialylated N-glycans with colors corresponding to sialylation type. The chromatograms showed a comprehensive look with the remarkable different glycan compositions and structures between two groups, particularly considering NeuAc or NeuGc sialylation patterns. Mono- and di-NeuGc sialylated glycans (sky blue and blue coded, respectively) were dominant in WT, while mono- and di-NeuAc sialylated glycans (orange and red coded, respectively) were prominently presented in DKO. WT showed NeuGc-sialylated complex/hybrid N-glycans with relative abundance of 64% and NeuAc-sialylated complex/hybrid N-glycans with relative abundance of 13%. On the contrary, DKO showed NeuAc-sialylated complex/ hybrid N-glycans with relative abundance of 73% and NeuGc-sialylated N-glycans with less than about 2% in spite of CMAH gene mutation. We confirmed the phenotype of Neu5Gc loss and Neu5Ac increase in DKO pigs with this MS result. Each of the identified glycan composition included two or more peaks corresponding to structural or linkage isomers (Figure 5c and 5d). The most frequent structures of sialylated N-glycan isomers identified by CID tandem MS were shown at *m/z* 868.98[M+3H]^3+^ for WT and *m/z* 693.59[M+3H]^3+^ for DKO, respectively. The MS aslo showed that di-NeuGc sialylated bisecting glycan was observed as a major component in WT, while mono-NeuAc sialylated biantennary glycan (*m/z* 693.59[M+3H]^3+^) was identified as the major molecule in DKO. Figure 6 revealed the comparison of relative abundances of N-glycans with different sialylation types between two groups. Interestingly, DKO showed increased neutral N-glycans and decreased of total sialylated complex/hybrid N-glycans, comparing to WT (*P*<0.001). Especially, it revealed significant decrease of di-sialylated glycans (*P*<0.05). Consequently, the statistical analysis revealed that the ratio for NeuAc linked N-glycans between WT and DKO was 1:5 whereas the ratio for NeuGc-linked N-glycans between WT and DKO was 37:1.

**Figure 5.**
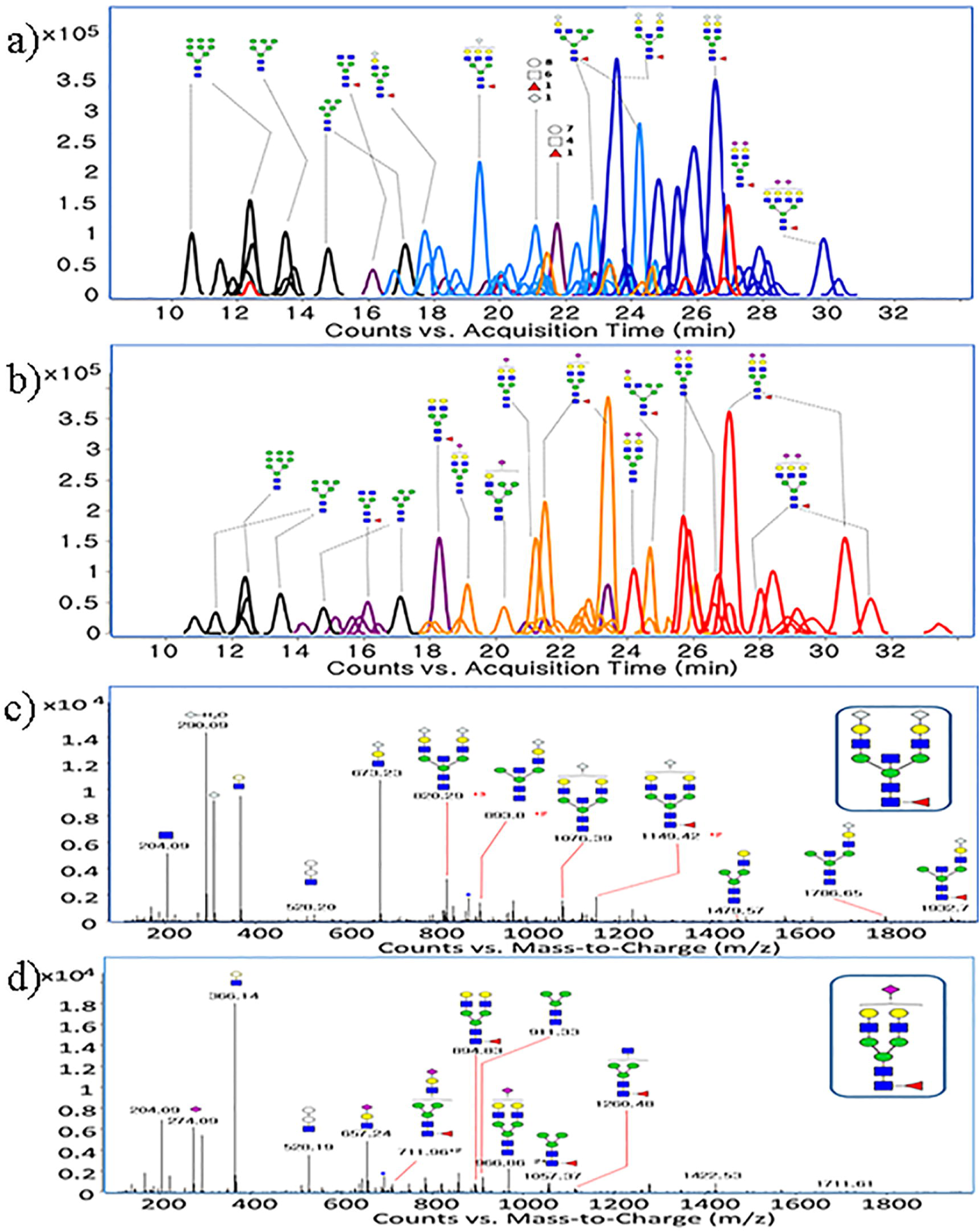
Comparison of N-glycan profilings obtained from erythrocyte membrane of WT and DKO pig. Extracted compound chromatiogram (ECC) of N-glycans of erythrocyte membrare from WT **(a)** and DKO**(b).** The ECCs were color coded according to sialylation types: black for high mannose type, purple for neutral type glycan, sky blue for mono-NeuGc sialylated glycan, blue for di-NeuGc sialylated glycan, orange for mono-NeuAc sialylated glycan, red for di-NeuAc sialylated glycan. Representative CID MS/MS spectra of major N-glycan with over 15% normalized intensity in WT**(c)** and major N-glycan with over 20% normalized intensity in DKO**(d)**

**Figure 6.**
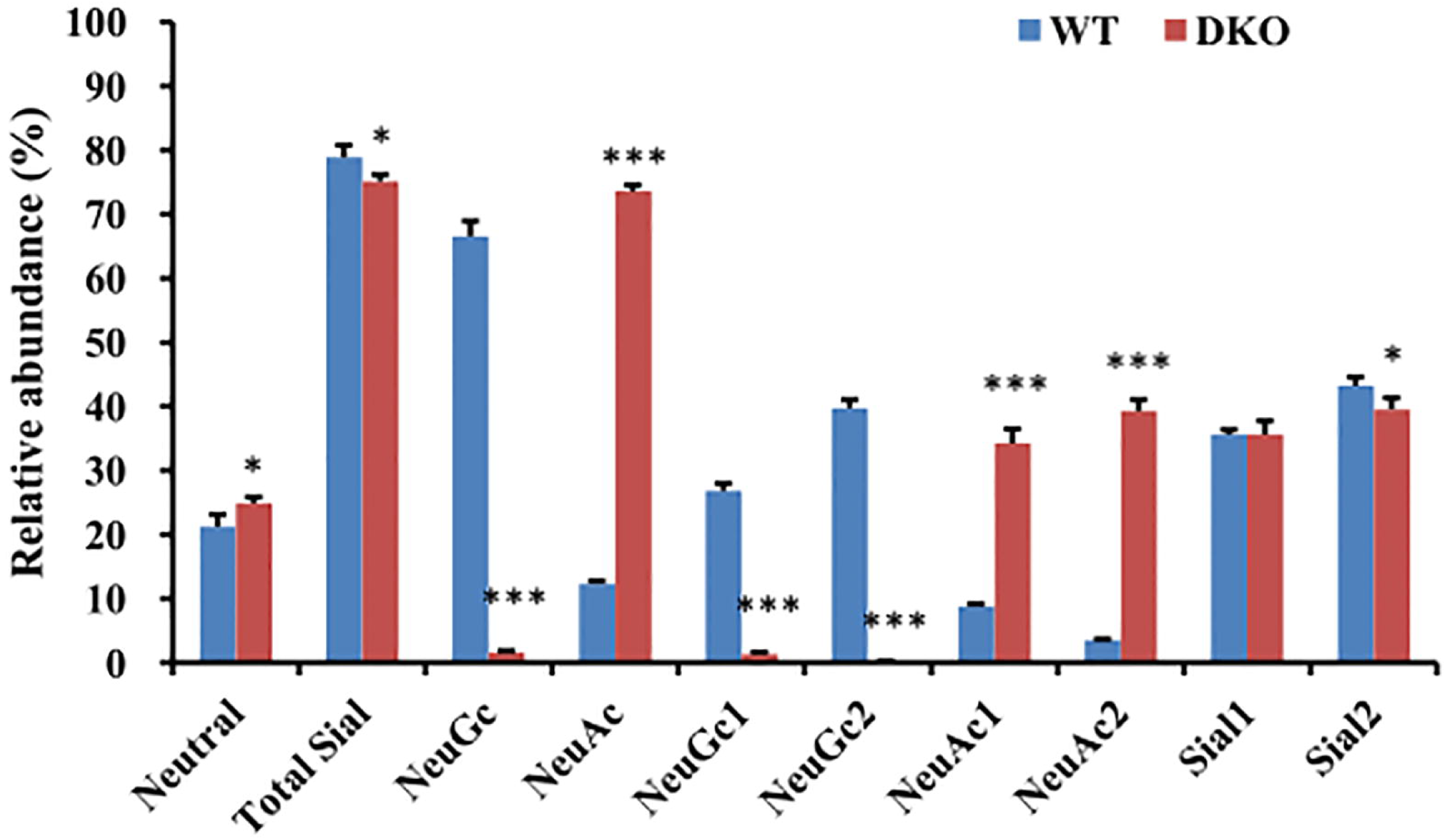
Relative quantitative comparisons of N-linked sialylated and neutral glycans. Total sial: NeuAc+NeuGc, NeuGc: mono-NeuGc + di-NeuGc, NeuAc: mono-NeuAc + di-NeuAc, Sia 1: mono-NeuAc+ mono-NeuGc, Sia 2: di-NeuAc + di-NeuGc, NeuGc1: mono-NeuGc, NeuGc2: di-NeuGc, NeuAc1: mono-NeuAc, NeuAc2: di-NeuAc (***P value <0.001, *P value < 0.05; P values were derived from the two-tailed Student t-test. Error bars show SEM)

### Lectin blotting with porcine erythrocyte membrane proteins

The coomassie blue staining followed after SDS-PAGE showed no difference in total protein expression levels between WT and DKO pigs (Figure 7a). In order to identify the glycosylation changes, we used seven kinds of lectins such as AAL (Fucα1-6GlcNAC,Fucα1-3(Galb1-4)GlcNAC), AOL (Fucα1-6GlcNAC(coreFuc)), UEA (Fucα1-2Galβ1-4GlcNAc) and LCA (Fucα1-6GlcNAc (core GlcNAc of N-linked glycopeptides)) for fucosylation and MAH (Siaα2-3Galβ1-3GalNAc), MAL (Siaα2,3Galβ1-4GlcNAc) and SNA (Siaα2-6Gal/GalNAc) for sialylation. The lectin binding assays of AOL (Figure 7b), AAL (Figure 7c), LCA (Figure 7d), UEA (Figure 7e), SNA (Figure 7f), MAH (Figure 7g) and MAL (Figure 7h) showed lectin binding reactivity with RBC membrane glycoproteins. The binding affinity signals with UEA and MAH showed no significant difference between the two groups, while the binding affinity signals with AAL, AOL and LCA lectins showed significant difference. It indicated the decrease of both terminal fucosylation and core fucosylation in DKO when compared to WT. Interestingly, MAL lectin showed the increased binding signals, suggesting the increased expression of terminal α2,3 sialic acid with the structure of Siaα2,3Galβ1-4GlcNAc.

**Figure 7.**
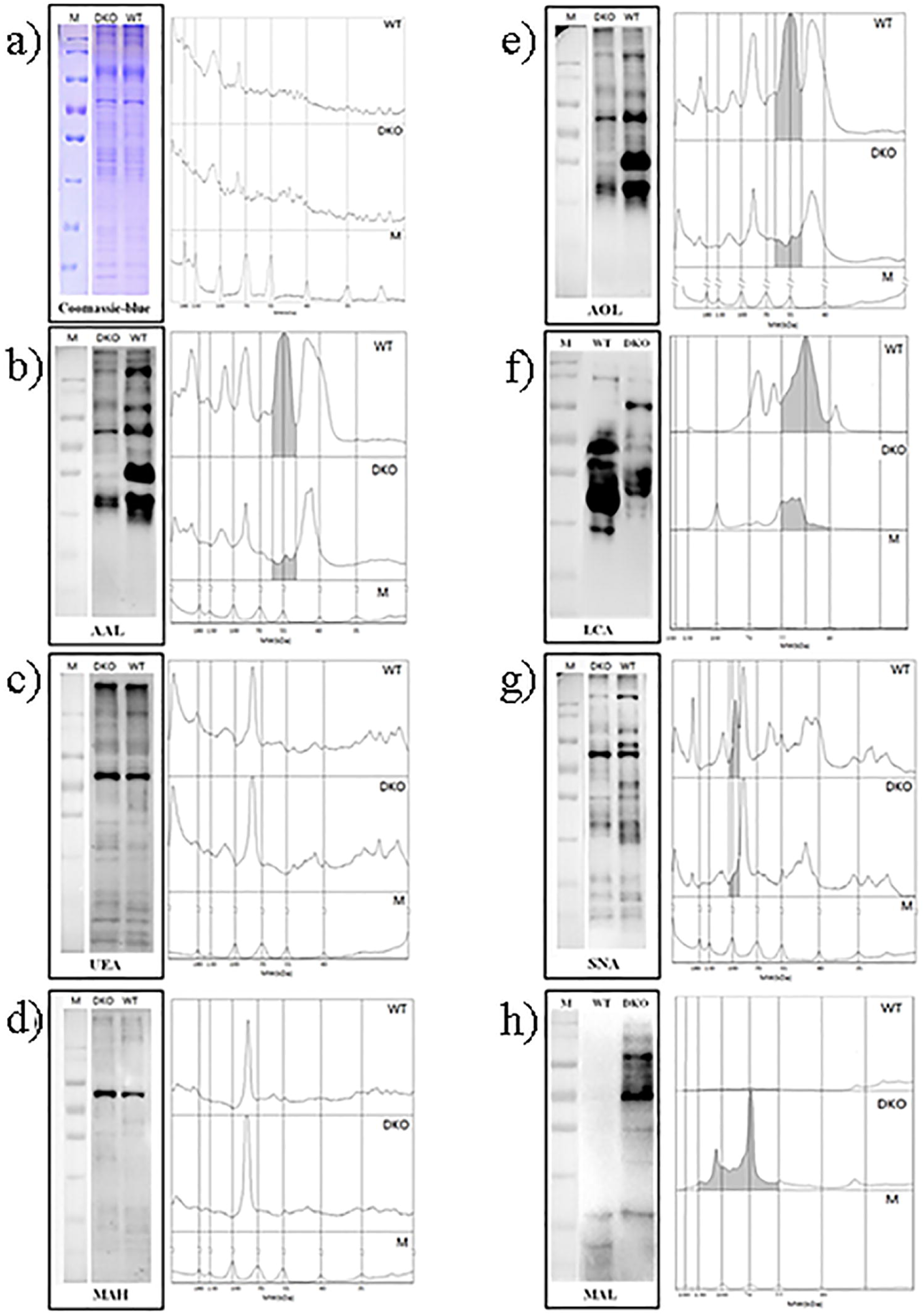
SDS-PAGE coomassie blue staining and lectin blotting analysis of GGTA1/CMAH double KO and WT pig erythrocyte membrane proteins. Double KO and WT **(a)** stained with Coomassie brilliant blue, **(b)** stained with AAL lectin, **(c)** stained with AOL lectin, **(d)** stained with LCA-lecin, **(e)** stained with UEA-I lectin, **(f)** stained with SNA lectin, **(g)** stained with MAH lectin and **(h)** stained with MAL lectin. The lectin binding intensity peaks from Image J software are shown on the right. Numbers on the right bottom indicate the molecular mass standards (kDa). M (molecular marker); WT (wild type); DKO (GGTA1/CMAH double KO)

## DISCUSSION

Pigs are considered potential donors for human transfusion and the recent progress of genetically modified pigs suggest that RBCs from pigs may eventually be able to be used for blood transfusion in humans(Long et al. 2010). The creation of GGTA1/CMAH gene double knockout in pigs has made it possible to consider the clinical application of pig erythrocytes for transfusion(Roux et al. 2010; Hui et al. 2007). However, GGTA1/CMAH DKO pigs has reduced but not eliminated the impact of human antibody binding and cell lysis in vitro(Lutz et al. 2013). The study on serum protein of GGTA1/CMAH DKO pigs showed that the inactivation of enzymes involving in carbohydrate modifications changed glycosylation patterns or created new potential carbohydrate xenoantigens which are not detected in humans and domestic pigs(Burlak et al. 2013). We produced GGTA1/CMAH double mutated piglets through CRISPR/Cas 9 and enrichment reporter system with eGFP and H-2K^k^, requiring three round SCNT. Our results showed that GGTA1/CMAH DKO pigs changed the sialylation pattern of N-glycans on the erythrocyte membrane as well as total terminal sialylation and fucosylation profiling on the erythrocyte membrane glycoproteins.

In this study, the immune reactivity of the DKO erythrocytes with human blood type O serum was remarkably decreased comparing to WT in hemagglutination assay. This result showed no discrepancy from the past study and supported the fact that DKO pig erythrocytes significantly reduced the agglutination comparing to GGTA1 KO and WT(Wang et al. 2014).

MS analysis in Figure 5a and 5b showed comprehensive chromatogram peaks representing N-glycans with different terminal sialic acid types (Neu5Gc and Neu5Ac) between WT and DKO. It was clear that in DKO, Neu5Ac was predominant whereas in WT, Neu5Gc was predominant. Interestingly, our result identified that less than 2% of Neu5Gc was still maintained in DKO. However, this was not surprising because it was reported that Neu5Gc was detected in human and DKO pigs(Burlak et al. 2013; Tangvoranuntakul et al. 2003). The Neu5Gc of DKO piglet RBCs in this study can be absorbed onto N-linked glycans from dietary sources. Further analysis is needed to clarify whether dietary incorporation of Neu5Gc explains the terminal sialic acid on DKO piglet RBCs. Moreover, Figure 5c and 5d revealed the CID tandem MS analysis to identify the most frequent glycan structures among sialylated N-glycan isomers between two groups. WT showed bisecting di-sialylated N-glycans while DKO showed biantennary mono-sialylated N-glycans as a major component. Bisecting GlcNAc, the central branch of N-glycan expressed highly in brain and kidney, is biosynthesized by a glycosyltransferase GnT-III(Nishikawa et al. 1992). It was known that the bisecting GlcNAc was a general suppressor of terminal modification of N-glycans including fucose and sialic acid(Miyako et al. 2019). We suggest that glycosyltransferase GnT-III as the biosynthetic enzyme of bisecting GlcNAc in DKO may be suppressed by GGTA1 and CMAH gene mutation. Figure 6 indicated statistically that DKO increased the expression level of neutral type glycans and decreased the expression level of total sialylated complex/hybrid N-glycans comparing to WT pigs. The decrease may be related with the ratio of CMP-Neu5Ac and CMP-Neu5Gc. If CMP-Neu5Gc is not synthesized in the GGTA1/CMAH DKO pigs, the sialylthransferase (ST) that utilizes it as a donor substrate uses CMP-Neu5Ac rather than CMP-Neu5GC. Although STs can use both CMP-Neu5Gc and CMP-Neu5Ac as donor substrates, the priority between CMP-Neu5Gc and CMP-Neu5Ac is different among STs. ST6Gal 1 showed 4-7 times greater activity toward CMP-Neu5Gc than CMP-Neu5Ac, whereas there was no significant difference toward these two substrates in ST3Gal 1. Since porcine erythrocytes have CMP-Neu5Gc as the main sialic acid, the deficiency of CMP-Neu5Gc can decrease the terminal modification activity of STs and in the meanwhile, cause the incomplete terminal modification of the receptor substrates or the modification by other carbohydrates, finally resulting in the decrease of sialylated glycans and the increase of netural glycans. Especially, the removal of αGal and Neu5Gc in RBCs may reduce the terminal modification activity of the α2,6 ST and increase the ST3GAL4 enzyme activity. GGTA1/CMAH DKO may lead to the incomplete carbohydrates by the flow of substrate components used in the αGal and Neu5Gc synthesis, the changes in feedback control pathway and the deletion of the terminal modification.

Moreover, the lectin assays showed that the DKO erythrocyte membrane glycoproteins significantly decreased the fucosylation in 55-70 kDa protein fractions and changed the sialylation in 70-100 kDa protein fractions compared to WT. It is known that the protein fractions with 55-100 kDa in pig include Band 4.5, Band 4.2, Band 4.1, Glycophorin A and Band 3(Matei et al. 2004; Sharma et al. 2011). Although the function of glycophorin A, the most abundant sialoglycoprotein on erythrocytes, has not been fully elucidated yet, it mainly functions not only as a counterpart between erythrocytes and the external environment, but also as a loader of various blood group antigens(Aoki T. 2017). Glycophorin A binds to neutrophil siglec-9, a sialic acid-recognizing receptor known to alleviate the activation of innate immune cells, and the sialic acid-based “self-associated molecular pattern” on erythrocytes inhibits neutrophil activation in the blood stream(Burlak et al. 2005). Insufficient sialic acid epitopes may prevent inhibitory signaling by the siglec receptors, leading to the activation of the immune system followed by xenograft rejection(Lizcano et al. 2017). Interestingly, the sialic acids of porcine erythrocyte account for negative charges on the membrane surface and have a great influence on its mobility, shape, function and lifespan(Aminoff et al. 1976; Eylar et al. 1962; Huang et al. 2016). The sialylation changes on DKO pig erythrocyte membranes should be a matter to be studied for the chargeability and biological properties of erythrocyte membrane and the xenotransfusion in the future. Furthermore, for successful pig-to-human RBC xenotransfusion, in addition to GGTA1 and CMAH mutation, the genetic modifications should include the expression of the H antigen (to reduce human anti-nonGal binding)(Costa et al. 1999), CD46 or CD55 (to protect against complement-dependent cytotoxicity)(Diamond et al. 2001; Cozzi and White. 1995), CD47 (to inhibit antibody-independent and possibly antibody-dependent phagocytosis by macrophages)(Ide et al. 2007) and CTLA4-Ig (to suppress T-cell activation and thus prevent sensitization and the production of anti-pig elicited antibody)(Vaughan et al. 2000; Hara et al. 2011).

In conclusion, the removal of αGal and Neu5Gc in pigs decreased the total fucosylation and sialylated N-glycans on the erythrocyte membrane. The present results will support further investigation into glycosylation alternation on GGTA1/CMAH DKO porcine erythrocytes and contribute to uncover the roles of glycan changes for clinical xenotransfusion.

## MATERIALS AND METHODS

### Animals

The oocytes were collected from unknown pigs at a local slaughterhouse. The nuclear donor cell lines were obtained from ear fibroblast cells of a 6-month-old female miniature pig. This research was carried out in strict accordance with the guidelines for the care and use of animals of Yanbian University. All animal experimental procedures were approved by the Committee on the Ethics of Animal Experiments in Yanbian University, Jilin, China (Approval ID: 20130310).

### CRISPR/Cas9 construction and enrichment of cells with mutations

The CRISPR/Cas9 system used to induce DNA double-strand breaks in GGTA1 and CMAH locus was previously described(Cho et al. 2013). Cas9 protein was complexed with CRISPR RNA (crRNA) and trans-activating crRNA (tracrRNA), forming a sequence-specific endonuclease that cleaves foreign genetic sequences. A single-chain chimeric RNA produced by fusing crRNA and tracrRNA replaced the two RNAs in the Cas9-RNA complex, resulting in a single-guide-RNA:Cas9 endonuclease (sgRNA:Cas9). The sgRNA:Cas9 induced site-specific genome modification and the target site was selected based on microhomology patterns to introduce frame shifting mutations at a high frequency(Bae et al. 2014). To get efficient enrichment of gene-modified cells, we employed the reporter vector construction including surrogate reporter (eGFP) and the magnetic reporter (H-2K^k^)(Kim et al. 2011; Kim et al 2013). The mouse H-2K^k^ gene was amplified from pMACS K^k^ (MiltenyiBiotech, Germany) and cloned into modified pRGS vector by isothermal cloning.

### Cell culture, transfection and enrichment of cells with mutations

The porcine ear fibroblast cells were obtained from of a 6-month-old blood type A female miniature pig to produce nuclear donor cell lines. The porcine ear fibroblasts were cultured and transfected as previously described(Kang et al. 2017). The fibroblasts were cultured for four passages in DMEM (HyClone) supplemented with 1% non-essential amino acids, 100 U/mL penicillin, 100 mg/mL streptomycin, and 15% FBS. The plasmids consisting of Cas9 and single guide RNA (total 10 μg, a weight ratio of 1:3, respectively) were transfected into the aliquots of 1× 10^6^ porcine primary ear fibroblast cells by electroporation. The transfected cells were cultured for 48 hours at 37°C and subjected to magnetic separation. Briefly, trypsinized cell suspensions were incubated with magnetic bead-conjugated antibody against a surface antigen, H-2K^k^ (MACSelect K^k^ microbeads; Miltenyi Biotech, Cologne, Germany), for 15 minutes at 4°C. Labeled cells were separated using a column (MACS LS column; Miltenyi Biotech) according to the manufacturer’s protocol.^24^

### Generation of fetuses and offspring by SCNT(Somatic Cell Nuclear Transfer)

H-2k^k^-positive cells with GGTA1 gene mutation were cultured for two additional days and were used as donor cells in first round SCNT. SCNT was performed as previously described(Yin et al. 2002). Oocytes were collected from unknown pigs at a local slaughterhouse. The mature eggs with the first polar body were cultured for 1 hour in medium supplemented with 0.4 mg/mL demecolcine and 0.05 M sucrose. Sucrose was used to enlarge the perivitelline space. Treated eggs with a protruding membrane were transferred to medium containing 5 mg/mL cytochalasin B and 0.4 mg/mL demecolcine, and the protrusions were removed with a beveled pipette. A single donor fibroblast was injected into the perivitelline space of each egg, followed by electrical fusion using two direct current pulses of 150 V/ mm for 50 μs each in 0.28 M mannitol supplemented with 0.1 mM MgSO_4_ and 0.01% polyvinyl alcohol. Fused eggs were cultured in NCSU-37 medium for 1 hour. The eggs were subsequently cultured in medium supplemented with 5 mg/mL cytochalasin B for 4 hours, followed by activation via two direct current pulses of 100 V/ mm for 20 μs each in 0.28 M Mannitol supplemented with 0.1 mM MgSO_4_ and 0.05 mM CaCl_2_. Activated eggs were cultured in medium for 7 days in an atmosphere of 5% CO_2_ and 95% air at 39°C. After the first round of SCNT, cloned embryos at 2–4-cell stage after one day of culture were transferred to the oviducts of naturally cycling gilts on the first day of standing estrus. The recipient pigs were euthanized at Day 23 of gestation and the fetuses with homozygous in frame mutations in GGTA1 gene were selected by using T7E1 assays and sequencing. The fibroblasts from the homozygous GGTA1 mutated fetuses were transfected with CRISPR/Cas9 plasmids including CMAH mutation and H-2k^k^-positive cells with GGTA1/CMAH gene mutations were produced as previous. Two gene mutated H-2k^k^-positive cells were cultured for two additional days and were used as nuclear donor cells in second round SCNT. After the second round of SCNT, the fetuses were surgically collected on Day 26 of gestation and were analyzed for homozygous in frame mutations. The fibroblasts from the selected fetus were used for the third round of SCNT. The pregnancy was assessed ultrasonographically on Day 25. The cloned piglets were delivered naturally or by inducing labor via intramuscular injections of prostaglandin F2 alpha (Ningbo, China) on Day 113 of gestation. The piglets with no mutations also were cloned via same SCNT and named as WT cloned pig.

### Mismatch-sensitive T7E1 assay and sequencing

The T7 endonuclease 1 (T7E1) assay was performed as described previously(Kim et al. 2009). The genomic DNA was isolated using a DNeasy Blood & Tissue Kit (Qiagen, Hilden, Germany), according to the manufacturer’s instructions. The region of DNA containing the nuclease-targeting site was PCR-amplified. The amplicons were denatured by heating and annealed to form heteroduplex DNA, which was treated with 5 units of T7E1 (New England Biolabs, Ipswich,MA, USA) for 20 minutes at 37°C, and assessed by agarose gel electrophoresis. To confirm that the mutation had been introduced into the target allele, PCR amplicons spanning the target sites were purified using aGel Extraction Kit (Qiagen, Hilden, Germany) and cloned into the T-Blunt vector using a T-Blunt PCR Cloning Kit (SolGent, Daejeon, Korea). The cloned inserts were amplified and sequenced.

### FACS and immunofluorescence of αGal expression in pig skin fibroblasts

To evaluate αGal expression, FACS and immunofluorescence analysis were performed as previously described(Yin et al. 2010). The skin fibroblasts (1 × 10^5^ cells diluted in 200 μL PBS with 0.1% (wt/vol)) produced from DKO piglet were prepared and stained at 4°C for 45 minutes with Alexa 488-conjugated Griffonia simplicfolia isolectin IB4 (GS-IB4) (5 μg/mL, Invitrogen). The stained cells were washed twice and then analyzed by flow cytometer (BD Accuri™ C6 Plus). Thereafter, the skin fibroblasts (1 × 10^5^) cultured overnight with DMEM plus FBS on chamber slide were fixed with 95% alcohol for 30 minutes, and then incubated with FITC-conjugated GS-IB4 for 30 minutes on ice. Fluorescence was visualized with an epifluorescence microscope (Nikon, Tokyo, Japan).

### Blood serum and erythrocytes preparation

10 mL of blood samples was collected in heparinized vacuum tubes from WT cloned pig (blood type A), GGTA1/CMAH DKO pigs (blood type A) and healthy human male volunteer (blood type O). Pooled human serum was purchased from healthy human male volunteer (blood type O). Whole blood samples were centrifuged at 1500×g for 10 minutes at 4°C using Ficoll-Paque Plus for erythrocyte isolation. The isolated erythrocytes were washed three times with phosphate-buffered saline (PBS) and diluted 1:10 in PBS at room temperature.

### Hemagglutination assay

The assay was performed in 96-well round bottom plates as described previously(Long et al. 2010). Briefly, the erythrocytes from blood group A WT and GGTA1/CMAH DKO pigs and blood group O human were prepared and suspended at 2 × 10^7^/well. After washing with HBSS, the erythrocytes were incubated with a 2-fold serial dilution of heat-inactivated human serum (blood type O) in HBSS at 4°C overnight. The images were scanned with an epifluorescence microscope (Nikon, Tokyo, Japan).

### Erythrocyte membrane purification

Erythrocyte membranes were prepared as previously described(Hanahan and Ekholm. 1974; Aoki et al. 2011; Bladier et al. 1979). 1 mL of packed RBC(red blood cell)s was lysed by mixing with 40 volumes of cold 0.01M Tris-buffer, pH 7.4 with 1mM phenyl methyl sulphonyl fluoride (PMSF), votexing per 15 minutes for 4 hours. The suspension was centrifuged with 13,000×g at 4°C for 20 minutes. The red-colored supernatants were discarded. The membrane pellets were re-suspended with 0.01M Tris-HCl buffer, pH 7.4 and centrifuged in the same manner as above. With the repeat re-suspensions, the membranes were washed four times until milky white preparation. After the final wash, the membranes were washed three times in cold PBS with PMSF and then stored at −80°C until the next analysis.

### N-glycan analysis by ECC(Extracted Compound Chromatogram)s and CID MS/MS

The N-linked glycans were prepared as previously described(Hua et al. 2011). The N-glycans were isolated from 100 μL of RBC membranes (concentration, 1.8 μg/μL) diluted with PBS/PMSF through in-gel digestion and peptide N-glycosidase F (PNGase F) treatment. The isolated N-glycans were purified and enriched through PGC-SPE (solid phase extraction) and detergent removal column, and were analyzed by NanoLC-Q-TOF-MS system. Glycan compositions were identified based on the newly established *in-silico* N-glycan library for pig specific glycans using mass accuracy and the LC retention time (RT). Each glycan quantity was normalized using the total abundances of all observed glycans. Additionally, the most frequent glycan structure among glycan isomers was elucidated by collision–induced dissociation tandem MS (CID tandem MS).

### SDS-PAGE and lectin blotting with porcine erythrocyte membrane proteins

Sodium dodecyl sulfate polyacrylamide gel electrophoresis (SDS-PAGE) was performed according to Laemmli with some modifications(Laemmli. 1970). Purified erythrocyte membranes were solubilized with ice-cold RIPA lysis buffer (strong type, P0013B, Beyotime) containing phenylmethylsulfonyl fluoride (PMSF) (P0100, Solarbio®) for 40 minutes on ice and centrifuged at 13,000×g for 15 minutes at 4°C. The supernatant was collected and the concentration of proteins was measured by BCA(bicinchoninic acid) assay using bovine serum albumin (BSA) as a standard (BCA Protein Assay Kit, Thermo Fisher, 23227). Sample was mixed in loading buffer (P1016, Solarbio®), heated at 100°C for 15 minutes and loaded on gels. Proteins were separated by SDS-PAGE in Bio-Rad mini-protean® Tetra system using PageRuler prestained protein ladder (#26616, Thermo scientific) as molecular-mass standards. Slab gel (1.5 mm thickness) consisted of 10% acrylamide in running gel (pH 8.8) and 5% acrylamide in staking gel (pH 6.8). After electrophoresis, gels were removed from the glass plates and were stained for protein by rocking for 1 hour at room temperature in a solution containing 0.5% w/v Coomassie Brilliant Blue R250 (Sigma), 45% v/v methanol and 10% v/v acetic acid. Gels were destained by constant agitation in several changes of 45% v/v methanol, 10% v/v acetic acid until maximum contrast was achieved between the stained bands and the background. After brief washing in water, the gels were photographed. For lectin-blotting, the separated proteins by SDS-PAGE were transferred to polyvinylidene difluoride (PVDF) membrane (EMD Millipore), washed with TBST for 5 minutes and then were blocked with 5% BSA in TBST[0.1%(v/v) Tween20-containing TBS, pH 7.4]. After blocking, PVDF membranes were incubated overnight at 4°C with AOL-Biotin Conjugate (A2659, TCI), biotinylated AAL (B-1395-1, VectorLabs), biotinylated LCA (B-1045, VectorLabs), biotinylated SNA(B-1305-2, VectorLabs), biotinylated UEA I(B-1065-2, VectorLabs), biotinylated MAL (B-1315, VectorLabs) and biotinylated MAH(B-1265, VectorLabs) (dilution 1:1000-2000 in TBST), separately. The PVDF membranes were washed six times each 5 minutes with TBST and then incubated for 1 hour at room temperature with HRP-labeled Streptavidin (A0303, Beyotime) diluted 1:5000 in TBST followed by washing as above. Bands were visualized by ECL Western Blotting Substrate (Thermo Fisher, 32209) and analyzed using ChemiDocTM MP Imaging System & Image Lab software (Bio-Rad). The intensity values of bands were measured using Image J software.

### Statistics

Each experiment was repeated at least three times. Data are presented as the mean±SD and are compared using a Student’s t-test. *P* values < 0.05 are considered statistically significant.

## Acknowledgement

The authors would like to thank our colleagues in the Jilin Provincial Key Laboratory of Transgenic Animal and Embryo Engineering.

## Author contributions

Xi-Jun Yin and Hyun Joo An designed the experiment. Choe wrote the manuscript. Experiments were done and discussed by all authors. All authors approved of the final manuscript.

